# “Spurring and Siloing: Identity Navigation in Scientific Writing Among Asian Early-Career Researchers”

**DOI:** 10.1101/2025.06.01.657048

**Authors:** Devon Goss, Meena Balgopal, Shaila Sachdev, Grace Kim, LaTonia Taliaferro-Smith, Sarah C. Fankhauser

## Abstract

Scientific writing and publication are critical sites where students must navigate between personal identities and presumed scientific objectivity. For non-white student researchers, particularly those stereotyped as “model minorities,” this process involves complex negotiations. This qualitative study examines how Asian and Asian American students navigate ethnic and cultural identities within scientific writing processes. Using an integrated theoretical framework combining cultural community wealth, narrative identity, and border crossing, we analyzed how participants deployed cultural resources while navigating institutional expectations. Drawing on interviews with 23 student participants who engaged with the [Blinded Journal], we identified two primary approaches: “spurring,” wherein ethnic and cultural backgrounds catalyze research questions, and “siloing,” wherein these identities are deliberately compartmentalized during writing. Findings reveal that while participants drew upon familial capital to inform research interests, most practiced disciplinary compartmentalization and adhered to perceived scientific norms. This tension demonstrates how students negotiate cultural scripts while developing professional identities. Our findings suggest that science educators could help students recognize their cultural backgrounds as valuable epistemic resources rather than sources of bias, explicitly address the myth of value-neutral science, and provide models of how scientists integrate personal and cultural perspectives into rigorous scientific work.

## Introduction

Science is inseparable from the act of communicating science: without communication, scientific knowledge cannot exist, circulate, or advance. As students build science disciplinary literacies, they develop disciplinary knowledge (Shanahan & Shanahan, 2008). Through engaging in science reading, writing, and speaking activities, for example, they gain a deeper understanding of how scientific knowledge is generated and evaluated by peers (Norris & Phillips, 2003). Subsequently, if given opportunities to engage in academic science writing, students learn what is valued in the science community (Sampson et al., 2013; Authors 2021; 2022). For this reason, scientific writing and publication represent critical junctures in science education when student researchers develop academic and professional identities as well as learn to navigate the cultural norms of their academic disciplines (Cameron et al., 2020; McRell et al., 2021; Richardson, et al., 2021). For adolescents, who are already exploring the many facets of their emerging identities, developing a science identity through academic writing may or may not be challenging (Knain, 2013; Reveles et al., 2004).

For culturally and linguistically diverse (CLD) students, the academic writing process may involve complex negotiations between their identities and the presumed objectivity of scientific discourse taught in academic settings. These negotiations are particularly salient in the natural sciences, where dominant epistemological frameworks emphasize universal knowledge and researcher neutrality (Yore et al., 2004). However, students may be empowered by drawing on their personal cultural and linguistic capital, discourse practices, and epistemic stances through translanguaging (Licona & Kelly, 2020). Translanguaging, the phenomenon of seamlessly integrating diverse communication norms to make meaning, is distinct from code-switching in that translanguaging occurs when people draw on multilingual knowledge to make meaning of concepts (Karlsson et al., 2019). Teachers use translanguaging pedagogical approaches when they aim to empower students to learn new concepts using any funds of knowledge meaningful to learners (Henderson et al., 2021; Pun & Tai, 2021). However, in science writing activities that are co-curricular and outside of classrooms, it is less clear how students draw on these various capitals necessitating a study on how CLD science students engage in science writing outside the classroom.

In this paper, we examine how Asian and Asian American science students negotiate their identity as they engage in scientific writing and publishing. Specifically, we sought to identify whether and how participants draw on resources and cultural and linguistic capitals as they wrote and published their scientific papers. This paper has been posted as a preprint (Authors, 2025).

### Research Gap and Significance

While substantial research has examined barriers facing minoritized groups in science, less attention has been paid to how science students actively construct academic identities that integrate or compartmentalize their cultural and linguistic backgrounds. This gap is particularly significant given growing evidence that cultural resources and perspectives can enhance scientific inquiry (Medin & Bang, 2014; Hogg & Volman, 2020) while simultaneously creating tensions with dominant scientific norms (Hsin & Xie, 2014). Understanding how students navigate these dynamics is crucial for developing science education practices that support diverse learners and prepare inclusive scientific practitioners and for supporting the development of students who pursue research experiences (Licona & Kelly, 2020).

Recent research on reflexivity highlights the unavoidable influence of researchers’ social locations—their cultural identities, institutional affiliations, and relationship to power—on scientific practice (Oswald, 2024; Reich, 2021). Despite this, the integration of personal identity into scientific work remains contentious, with some scholars arguing that acknowledging positionality violates scientific principles of universalism (Savolainen et al., 2023). This tension presents potential challenges for science students from non-dominant backgrounds as they develop their scientific voices and academic identities in a space that often excludes explicit incorporation of one’s identity. Here, we use non-dominant to mean students who may be less likely to encounter authors of scientific papers who share the same ethnic/cultural identity, but who study in schools with peers and teachers who do share their same ethnic/cultural identity. While some scholars focus on tensions, others focus on the assets that students from non-dominant backgrounds bring to academic discourse, calling out opportunities for students to integrate different funds of knowledge, such as through translanguaging (Lemmi et al., 2022). However, much of the literature on translanguaging in STEM education focuses on students who identify as African American or Latine (e.g., Brown, 2021) and often are conducted in classroom settings analyzing oral discourse (e.g., Licona & Kelly, 2020; Pun & Tai, 2021; Rafi & Morgan, 2023). Because writing is a form of meaning making, understanding how students draw on their suite of capitals to succeed in science studies and inquiry is important. There is a need, therefore, to examine the experiences of students who identify as Asian or Asian American when engaged in academic science writing.

### Asian and Asian American Scientists as Critical Case

The experiences of Asian and Asian American students offer particularly rich insights into the dynamics of reconciling personal and professional ways of communicating science. The model minority stereotype frames Asian and Asian Americans as naturally skilled in science and mathematics while simultaneously marking them as perpetual foreigners in scientific institutions (Lee, 1996; Chinn, 2002; Steele, 2011). This creates both unique pressures and opportunities as these science students navigate cultural expectations, professional norms, and ethnic stereotypes. Recent research suggests that Asian American academic achievement stems primarily from cultural values around effort and immigration-related factors rather than individuals’ innate abilities or advantage (Hsin & Xie, 2014). These findings challenge simplistic narratives while highlighting how cultural values persist across generations, shaping how Asian and Asian American scientists approach their work. However, the model minority framework simultaneously masks diversity within Asian and Asian American experiences while creating additional pressures through unrealistic achievement expectations and limited identity expressions (Hsieh & Kim, 2020; Poon et al., 2016).

## Theoretical Framing

To understand how Asian and Asian American science students navigate identity in scientific writing, we integrate several theoretical perspectives that together illuminate both individual agency and institutional constraints. These theories -- cultural community wealth, narrative identity model minority discourse, and border crossing -- complement each other by revealing different ways that individuals draw upon cultural capitals, construct meaningful identities, and navigate institutional pressures in scientific contexts. Collectively, these three frameworks illuminate how science students may engage in border crossing from cultural to academic spaces and back again Rios-Aguilar & Neri, 2023; Esteban-Guitart, 2016).

### Cultural Community Wealth in Scientific Practice

Yosso’s (2005) cultural community wealth (CCW) framework illuminates how non-white scientists may draw upon interconnected forms of community-nurtured resources to enhance scientific inquiry and create pathways for innovation. These capitals are diverse and include family (social support), aspirational (maintaining a positive outlook), linguistic (ability to leverage different languages), navigational (accessing resources to reach academic or professional goals in challenging environments), and resistant capitals (challenging barriers for success). Rather than viewing cultural background as a deficit to overcome, the CCW framework reveals how individuals draw on different forms of capital that might inform their research questions, help them develop methodologies, and contribute to scientific knowledge. The process often begins with scientists drawing on familial and community knowledge to identify research questions that mainstream science might overlook. For example, Kimmerer (2015) documented how Indigenous scientists integrate traditional ecological knowledge passed down through generations into their research, leading to innovative approaches in environmental research that challenge and enrich “Western” scientific paradigms. Like Rodriguez and colleagues (2023) found that women computer science students drew on family members’ knowledge of the computer science workforce and culture, regardless of whether they had degrees in computer science or not. In other words, CCW assumes that family and community capital are an asset for science students to draw upon. It also may provide opportunities for familial capital to inspire students’ initial research questions and provide alternative frameworks for data interpretation and analysis.

Community connections extend beyond knowledge systems to create practical pathways for research impact. Zambrana and colleagues (2015) described how scientists from minoritized groups leverage social capital through community networks to develop research partnerships and access study populations, while simultaneously building trust between scientific institutions and communities. The trust-building process is crucial for conducting more comprehensive research while ensuring its relevance to community needs. Ethnic identity, alone, does not likely shape an adolescent’s academic identity; support from family and community members likely plays an important role (Wong, 2015). For example, Cultural Linguistic Identity (CLI), the integrated sense of self shaped by cultural heritage and linguistic repertoires, represents a foundational resource that scientists could bring to their work. When students grow up in multilingual families, it may expand their community connections and shape identity. For this reason, language plays a crucial role in this knowledge integration process, though not without complexity. However, as Bowker (2021) demonstrated, although multilingual abilities could theoretically enhance scientific communication and enable broader knowledge exchange, institutional pressures to publish in English often constrain scientists’ ability to fully leverage their linguistic resources. While multilingual scientists possess valuable capabilities for bridging different knowledge systems, they face structural barriers to deploying these capabilities effectively (Di Bitetti et al., 2017). This tension illustrates how CLI operates as both resource and constraint as multilingual scientists may be positioned as less credible when their linguistic repertoires diverge from monolingual English norms, even as these same repertoires could enrich scientific discourse.

The deployment of such cultural resources requires significant navigational skills, particularly in institutional contexts that may not readily recognize their value. Chinn’s (2002) work with Asian and Pacific Islander women scientists revealed how they drew on cultural traditions of adaptability and strategic relationship-building to successfully maneuver through male-dominated scientific spaces. Navigation is supported by aspirational capital, which may enable scientists to maintain their research vision despite institutional skepticism that has been notably documented towards women (Bird & Rhoton, 2021).

Perhaps most significantly, CCW enables scientists to develop resistant capital - the ability to question dominant paradigms and propose alternative perspectives. Scholars have demonstrated how Black women scientists leverage their critical awareness of power dynamics to identify gaps in scientific knowledge and develop new methodological approaches (Prescod-Weinstein, 2020; Rankin et al., 2021). Their resistance to limiting assumptions, supported by community knowledge and robust social networks, often leads to theoretical innovations that benefit the broader scientific enterprise. When the assumptions of scientists and science students are challenged, they discover that community knowledge can be essential capital supporting scientific discourse (Rodriguez et al., 2023; Weinberg et al., 2018). Together, these forms of capital form a rich foundation for scientific practice that extends beyond individual research capabilities to encompass community relationships, knowledge systems, and pathways for innovation (e.g., Authors et al, 2021). Understanding science students draw upon these resources - or face constraints in doing so - is crucial for developing more inclusive scientific institutions that benefit from diverse perspectives and approaches. CCW allows researchers to use an asset-based approach to understand the experiences of science students, but it does not focus on the entirety of experiences that an individual has. Here, narrative identity theory is relevant for our study.

### Narrative Identity and Cultural Scripts in Science

Narrative identity theory illuminates how scientists make meaning of their experiences through stories that are mediated by cultural symbolic systems (Ricoeur, 1991). For science students from non-dominant backgrounds, these narratives bridge multiple cultural worlds: their home cultures, academic science culture, and broader societal narratives about race and achievement. The stories scientists tell about themselves—who they are, what knowledge they possess, and how they came to do science—are constrained and enabled by available cultural scripts.

Cultural scripts powerfully shape both how individuals see themselves and how institutions treat them (Chinn, 2002). In science, professional norms emphasize objectivity, detachment, universal knowledge, and individual achievement. These norms often conflict with cultural scripts that emphasize community obligations, cultural knowledge transmission, collective achievement, and specific forms of identity expression. Scientists from non-dominant backgrounds must navigate between competing narrative possibilities: they can construct professional identities that align with dominant scientific scripts (potentially requiring erasure of cultural identity), or they can attempt to construct narratives that integrate cultural identity into scientific work (risking being marked as non-credible or biased). For instance, Medin and Bang (2014) documented how Indigenous scientists must reconcile “Euro-centric” scientific emphasis on researcher neutrality with cultural traditions that acknowledge intimate connections between researchers and the phenomena they study. This represents a conflict between available narrative scripts: the dominant script positions researchers as objective observers outside the system, while Indigenous cultural scripts position researchers as participants within interconnected systems. The narrative identities Indigenous scientists construct must somehow navigate between these storylines about what it means to produce knowledge.

Language itself functions as both a medium and constraint for narrative identity construction in science. Hwang (2005) describes the dominance of English as the language of science, which thus presents a barrier to those on the “periphery” (i.e. outside Europe or English-speaking countries). The fact that scientists in the periphery must consume, digest, and produce scientific knowledge in English fortifies a power imbalance and reinforces competence in English as a key to scientific recognition. This linguistic requirement shapes the narratives that scientists can tell about themselves and their work. Multilingual scientists may possess rich scientific knowledge and sophisticated research insights, but if they cannot articulate these in fluent academic English, the narrative scripts available to them shifts from one of “expertise” to that of “deficiency.”

A notable example within the sciences that further highlights tension in identity within science production is the myth of the model minority. The model minority myth framework provides crucial context for understanding how institutional pressures shape identity navigation, particularly for Asian and Asian American scientists (Poon et al., 2016). This stereotype creates a complex web of expectations and constraints that influences how scientists construct their professional identities. Lee (1996) demonstrated how the model minority narrative simultaneously provides advantages through positive expectations about scientific ability while limiting acceptable forms of identity expression. This dynamic often requires Asian and Asian American scientists to carefully manage their self-presentation, potentially constraining their ability to bring cultural resources into their work (e.g., Nguyen et al., 2022). Interestingly, Archer and colleagues (2015) found that secondary science students identifying as South Asian or East Asian were more likely than those identifying as other ethnic groups to have high science capital through a survey. Subsequent work by this team found that the relationship between cultural and science capital was nuanced and not always correlated (Moote et al., 2021).

### Border crossing within academic spaces

Academic science has been described as a separate culture, with accepted norms of how to interact with others in the cultural community, to generate new information using accepted practices and protocols, and values of what is important to study. When students arrive in academic spaces bringing different worldviews other than those espoused in academic science classrooms, they may find acceptance of the new culture challenges (Aikenhead & Jegede, 1999).

Educators, therefore, may have to help students navigate across different epistemological spaces, so they can succeed in learning academic science (Aikenhead, 2001). For example, Borgerding (2017) described how a rural science teacher acted as a border crossing “tour guide” to help his students make meaning of both academic explanations for biological evolution and culturally relevant local funds of knowledge about the natural world. Without expressing judgement, the teacher allowed students to express cognitive dissonance as they learned new academic content, even if they did not end up accepting biological evolution as an explanation for biodiversity. Sanchez Tapia (2020), in another example, found that Indigenous students in a Mexican middle school science classroom crafted arguments that drew on both newly learned academic evidence and their own cultural knowledge. One of the participants, Carmen, believed that just as she benefitted from learning about academic epistemologies, that the scientific communities should benefit from learning about Nahua (Indigenous) ways of knowing. In other words, she saw the difference ways of knowing as complementary, as Author and colleagues (2021) reported in their research with Indigenous South Asian students enrolled in a biology course.

### Integration: A Multi-Level Framework

These three theoretical perspectives work together to illuminate different aspects of identity navigation in science. Cultural resources provide raw material for identity construction, while narrative processes transform these resources into coherent professional identities. Institutional contexts shaped by model minority myth condition individuals in how these resources can be deployed and how identities can be expressed. By border crossing, students must navigate how they move between spaces and decide how to deploy different funds of knowledge in each space.

This integrated framework allows us to examine both individual agency in drawing upon cultural resources, and institutional constraints on identity expression. It suggests that successful navigation requires both strategic deployment of cultural resources and careful attention to institutional demands. Moreover, it helps explain why scientists might simultaneously draw upon and distance themselves from cultural resources as they navigate different demands and expectations in scientific contexts.

### Research Questions

Building on this conceptual foundation, we developed three broad research questions to allow for an exploratory analysis of student experiences in professional scientific writing as they relate to the frameworks of CCW, narrative identity, and border crossing. Specifically, our study examines:

1. In what ways do Asian and Asian American science students draw upon different forms of cultural capital in their scientific research and writing processes?
2. How do Asian and Asian American science students view scientific writing as a space for identity integration?
3. How do Asian and Asian American science students navigate between scientific professional norms and their cultural backgrounds?

Using the critical case of Asian and Asian American science students to answer these questions, we aim to understand both individual strategies for identity navigation and broader implications for institutional change. Our findings reveal two primary approaches- which we term “spurring” and “siloing” - through which science students manage the relationship between their cultural identities and scientific work.

## Methodology

### Study Context and Significance

This survey and interview study focuses on science students publishing through the [Blinded Journal], a platform that supports high school students in developing research and scientific writing skills. The journal’s peer review process mirrors professional scientific publishing while providing supportive feedback from graduate student and postdoctoral reviewers. This context provides unique insight into how students navigate academic identity as they enter scientific publishing, as they are socialized into disciplinary norms. The findings contribute to understanding both individual identity navigation strategies and institutional factors that support or hinder identity integration in science.

### Participants and Recruitment

To examine how science students incorporate their cultural and linguistic identity (CLI) into scientific writing and publication, we conducted a comprehensive recruitment process through the [Blinded Journal]. We sent email invitations to 1,904 individuals who had previously participated in [Blinded Journal] as student authors, editors, or reviewers. From the pool of respondents, we selected potential participants based on several criteria: self-identification as a person of color, current involvement in scientific research or writing, either as a student or early-career scientist, and willingness to discuss their experiences with identity and scientific work in detail.

Of the initial respondents meeting our criteria, we contacted 45 individuals for interviews. In this paper, we analyze the interviews we conducted with twenty-three participants who completed interviews and self-reported their background to be of Asian or Asian American. These participants represent a range of ethnic and national backgrounds, scientific disciplines, and career stages (Table 1). Most participants (87%) were based in the United States, while 13% were international participants. More than half of the participants (56%) reported that they spoke a language other than English. This sample size aligns with recommendations for qualitative interview studies seeking to achieve theoretical saturation while maintaining depth of analysis (Charmaz, 2006). The study was approved by the [x] University Institutional Review Board [Study STUDY00000797]. All participants provided their informed consent or assent prior to completing the interview.

### Interview Protocol and Process

Our semi-structured interview protocol was designed to explore participants’ experiences based on our research questions, examining their personal and cultural background, early experiences with science and writing, current scientific writing practices, perceptions of identity in scientific contexts, experiences with peer review and publication, and navigation of cultural identity in scientific settings and scientific writing. Interviews were conducted virtually and lasted between 45-60 minutes each, generating approximately of recorded conversation. All interviews were audio-recorded with participant consent and professionally transcribed.

### Data Analysis

Our approach examines participants’ own narratives about how their CLI influences their scientific work, without assuming deterministic relationships between ethnicity and epistemology. We treat these self-reports as valuable data about identity navigation processes rather than evidence of epistemological differences. We therefore utilized a thematic analysis approach to this data. Analysis includes coding data in a multi-stage process aimed at understanding similarities and differences among the participants within their narratives￼￼.

Our thematic analysis approach manifested itself in two stages. In the initial collaborative coding phase, five members of the research team collectively analyzed one interview transcript to establish consistent coding practices and develop preliminary coding categories. Open coding was utilized to link phrases and ideas with each other into larger themes (Glaser, 1978). This process helped ensure reliability and allowed us to identify both anticipated themes from our research questions and unexpected patterns in the data. During this stage, the general codes of racial identity playing both a ‘spurring’ and a ‘siloing’ role in the narratives of scientific writing for the participants were uncovered. As these general codes were discovered, the authors viewed an additional subset of transcripts, leading to the development of the specific codes, as discussed in the next section. Through this collaborative process, we developed a codebook that included *a priori* codes derived from research questions and our theoretical framing, emergent codes identified through initial transcript analysis, clear definitions and examples for each code, and guidelines for code application. Three members of the research team then led the systematic coding of all transcripts. These researchers met weekly to compare coding decisions, discuss ambiguous cases, resolve disagreements through consensus, refine code definitions as needed, and document coding decisions. The first author continued to refine the codebook and reapply it across the transcripts.

### Positionality of the research team members

The research team varies in terms of their career stage and ethnic identities, which is helpful for gaining multiple perspectives on the study topic. Four of the authors are mid-career stage researchers. One of the authors identifies as a White American who only speaks English. One of the authors identifies as a White American who speaks only English. One of the authors identifies as a Black American who speaks only English. One of the authors identifies as South Asian American and familiar with three South Asian languages. Two of the authors are student researchers. One of the authors identifies as South Asian American and is bilingual in English and Spanish. One of the authors identifies as East Asian American and is bilingual in English and Korean.

## Findings

Our analysis revealed complex dynamics in how Asian and Asian American science students navigate CLI in scientific writing. When asked about how their Asian identities were incorporated into their process of writing and publishing a scientific article, the science students demonstrated the conflicting ways in which they approach integrating their ethnicity into their academic work. Two primary approaches emerged: spurring, wherein they cited their cultural and linguistic backgrounds and ethnicized experiences as important stimulation for their research and writing process; and siloing, wherein they stressed the fundamental need to separate CLI from the scientific writing and publishing process (Table 2). These strategies reflect different ways of leveraging cultural capital and navigating institutional expectations within scientific contexts.

### Spurring

When initially asked about the role that their race had played within their writing process, many of the Asian student authors that we interviewed indicated that they had not previously considered the role of their identity within their research process. However, upon further reflection, many indicated that they indeed felt that their CLI had influenced their experiences in academic publishing. Specifically, 39.1% viewed their Asian CLI as spurring the inquiry and framework for the project that led to their article publication. In so doing, these Asian student authors showcase the ways in which CLI can be used as a bridge between differing community values and scientific inquiry.

#### Familial Capital: Strong Evidence for Strategic Use

Our findings provide substantial evidence for how students draw upon their CLI in their scientific work, with familial capital serving as the foundation through which this cultural knowledge is accessed and deployed for scientific inquiry. This was particularly evident in how students leveraged cultural knowledge to inform their research choices and approaches.

Evan, a 17-year-old who identified as East Asian, exemplified this strategic use of familial capital when he reported why he felt motivated to ask questions about environmental issues in a Chinese context, stating,

> If I weren’t Chinese, maybe I wouldn’t have chosen the topic specifically, and maybe I wouldn’t have been so like concerned about an environmental injustice that might be happening. So, I think, being from this background, it kind of inspired me to choose this topic, and really like, resonate with the topic, and be more like, involved in research, and not do it for the sake of, you know, for the sake of doing it. It kind of made me more like passionate about like what I’m doing and made the process honestly a lot more easier, a lot easier because it gave me like motivation.

**Table 1:** Participant Demographics.

| **Self-Reported Demographic Information** | **Frequency** | **Percentage** |
| --- | --- | --- |
| **Self-Identified Ethnic Identity** |  |  |
| East Asian | 12 | 52.2% |
| South Asian or Indian | 6 | 26.1% |
| Southeast Asian | 3 | 13.0% |
| Central Asian | 1 | 4.3% |
| Pacific Islander | 1 | 4.3% |
| **Gender** |  |  |
| Male | 11 | 47.8% |
| Female | 12 | 52.2% |
| **Country of Residence** |  |  |
| USA | 20 | 87.0% |
| India | 1 | 4.3% |
| China | 1 | 4.3% |
| Vietnam | 1 | 4.3% |
| **Highest Education Level Completed** |  |  |
| Less than High School | 1 | 4.3% |
| High School | 6 | 26.1% |
| Undergraduate School | 10 | 43.5% |
| Graduate School | 5 | 21.7% |
| Not in School | 1 | 4.3% |
| **Science Discipline (Authors Only)** |  |  |
| Natural Sciences | 10 | 55.5% |
| A.I. & Data Sciences | 3 | 16.7% |
| Social Sciences | 3 | 16.7% |
| Environmental Sciences | 2 | 11.1% |

Note: $N=23$ participants

**Table 2.**
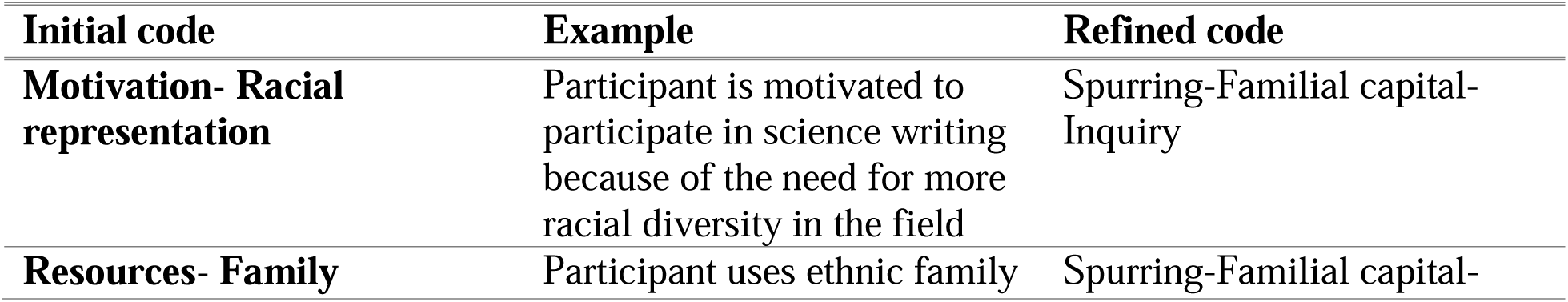
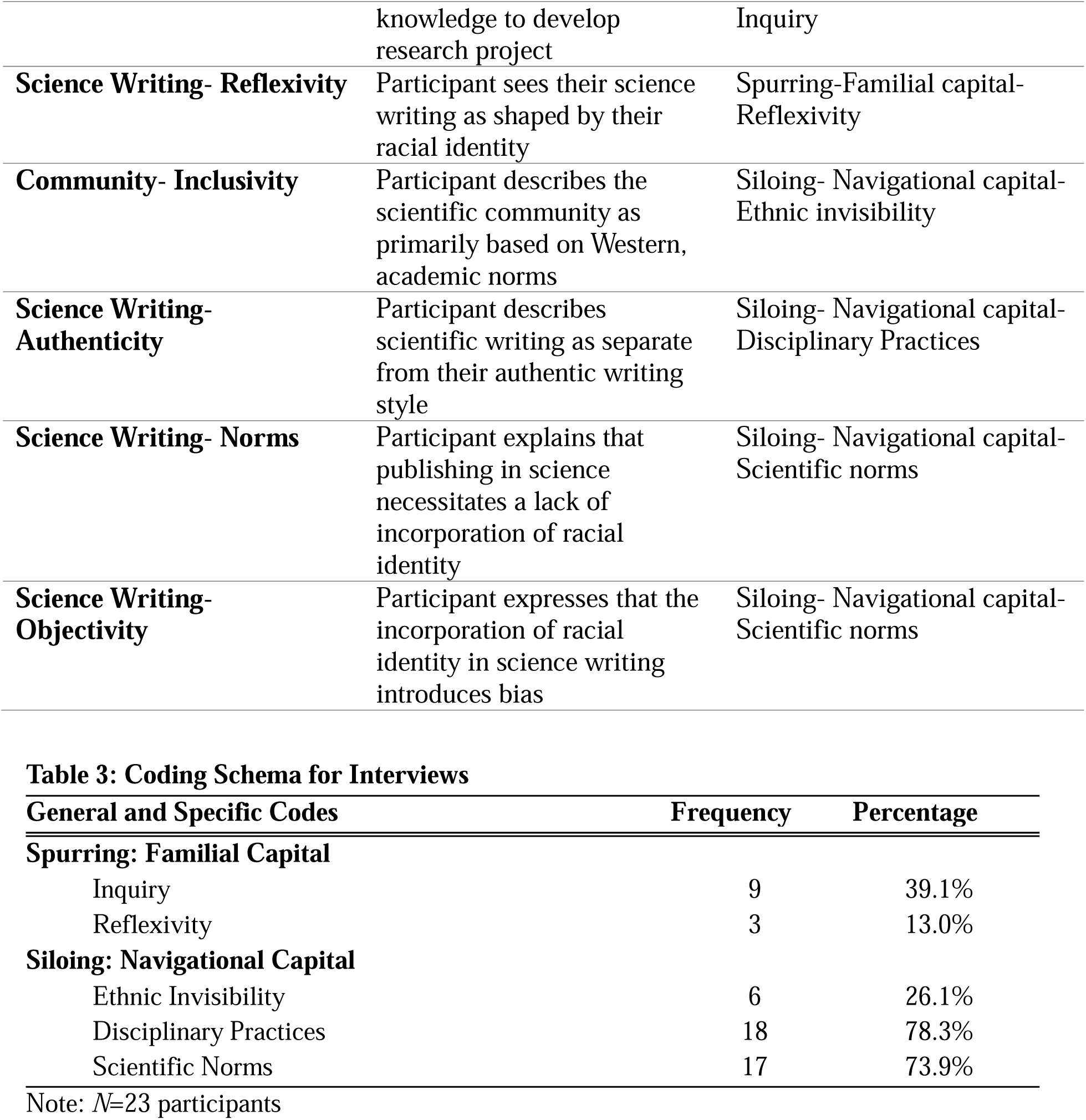
Emergence of the Theoretical Coding “Spurring” and “Siloing”.

**Table 3:**
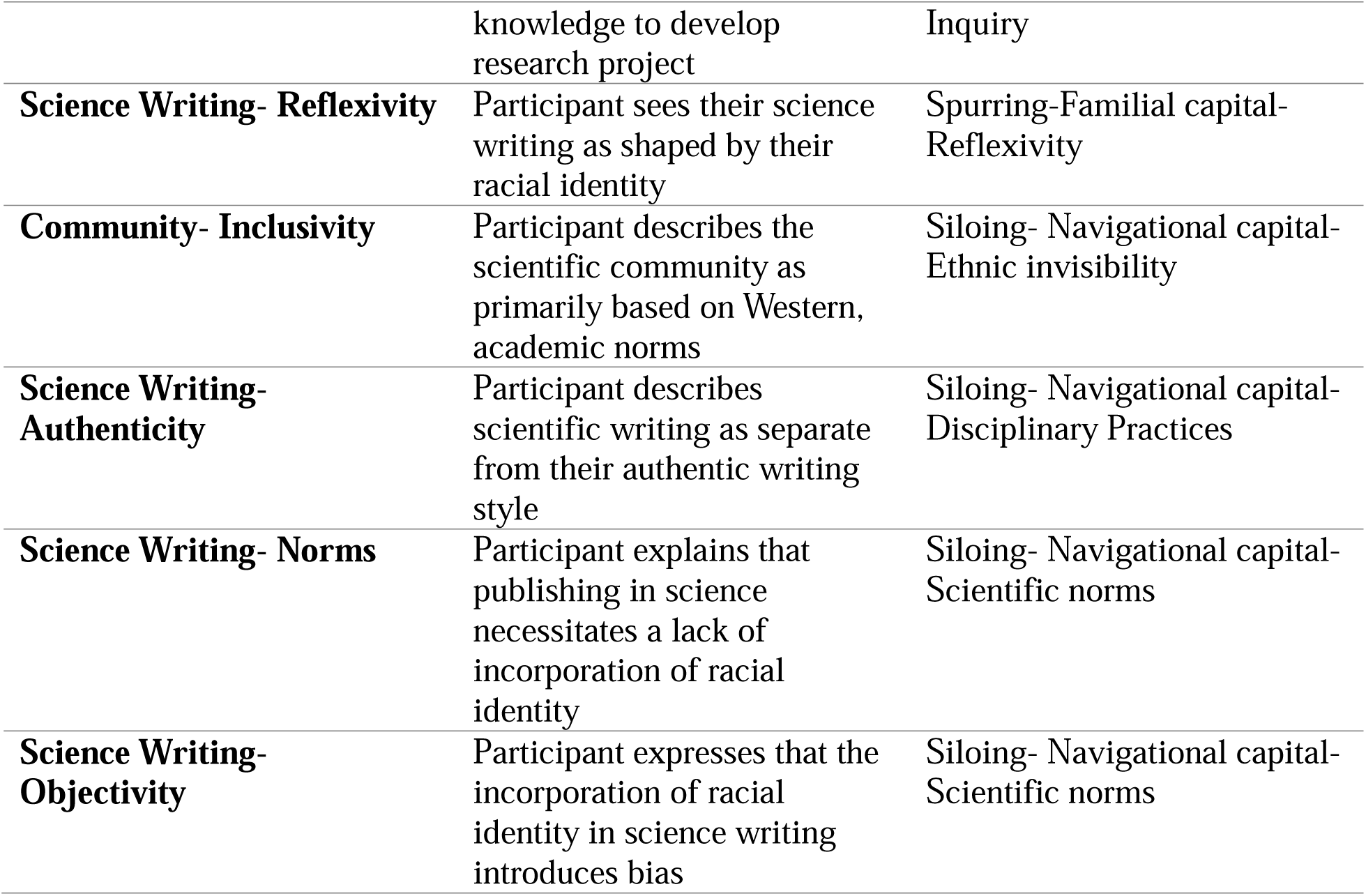
Coding Schema for Interviews.

His statement reveals how cultural knowledge transmitted through community connections shaped not just his topic selection but his sensitivity to environmental concerns. Similarly, Chloe, 18-year-old woman who identified as East Asian, shared how cultural knowledge can informed her scientific inquiry when she integrated traditional medicine into her teach research:

> I think I actually did mention [my culture] a little bit with my project, because mine was about tea. So, I kinda talked a little bit about like Asian, like traditional medicine, and like how tea is like has been used for a long time, in like maybe like 2 sentences. But it was still like a nice touch, and I felt connected.

As Chloe’s and Evan’s remarks demonstrate, their ethnic contexts and backgrounds served as exposure for several initial research questions that they brought into the academic setting. Using their familiar capital, Chloe and Evan were able to construct scientific questions that legitimized concerns in their ethnic communities.

For some participants, drawing on familial capital when developing a research question was also reported to be widespread among their peers in scientific spaces. Amithi, 18-year-old woman who identified as South Asian, stated,

> From what I’ve experienced at school like in my AP research class, when everyone was sharing their research ideas and like their topics like, I feel like everyone was connected to their topic in one way. Either it was usually their race…or, like other people brought their cultures into it….so I do see, like a lot of people bringing in their personal identity into their research.

This observation indicates a recognition, at least among students, that cultural knowledge can serve as a legitimate foundation for scientific inquiry.

Although not initially driven by identity-related questions, other Asian student researchers encountered racial/ethnic themes during their research that resonated with their own backgrounds. Aaron, a 17-year-old student who identified as both East Asian and white, started their project with a more general research question, but explained

> “From my literature review, I did see a lot about how minorities are affected by Alzheimer’s, and they have, like a 3 times higher mortality rate from Alzheimer’s, and that really affected me…So I think that if that was my initial inspiration for doing Alzheimer’s research, then.”

Very few of the students that we interviewed saw their research as being fundamentally shaped by their standpoint as an Asian. However, 13% did express the ways in which CLI influences how scientists approach their research and writing process. For example, Won Jun, a 29-year-old man who identified as East Asian, shared that “I don’t know, I think there are times when I can voice my opinion, and I guess in that sense, my opinions draw from my racial cultural background. I guess there’s that, so maybe like in a little in some ways.” And Chloe, the 18-year-old East Asian woman, expanded on this idea, saying that one’s ethnic background impacts “how they do data analysis, how they see a certain problem, um, how they interpret certain results.” This spurring approach reveals the narrative work that Asian and Asian American science students use to legitimize cultural knowledge within scientific frameworks. Through careful narrative framing, these student scholars can maintain their scientific credibility while still bringing their ethnic identities into their academic work. In this way, they are able to navigate between cultural and academic traditions. However, our data also reveal limitations in how explicitly this capital can be acknowledged in scientific writing. Chloe’s characterization of her cultural references as just “2 sentences” and “a nice touch” suggests a careful calibration of how much cultural context is appropriate to include. This measured approach reflects the constraints of scientific writing formats that may not readily accommodate extensive cultural contextualization.

### Siloing

In general, the further that the students were engaged in their research and publishing process, the less that the scholars felt that their Asian ethnic identities influenced their scholarship. More than a quarter (26.1%) of the participants had never considered the possibility of their ethnic identities shaping their participation in the scientific writing process. Aaron, who had before discussed how his East Asian and white identity had influenced his decision to study health inequalities in Alzheimer’s in the previous section, responded to a question about how he incorporated his identity into his writing by stating, “I mean it’s my writing, I guess, but that’s never really been something that I’ve thought about.” Similarly, Evan, who had expressed how his East Asian identity had spurred his interest in studying environmental justice issues in a Chinese community, said “Like, for me, those things of background of race and identity, they were never like a topic much really present in my life.”

Aaron and Evan internalized dominant scientific scripts which imply that scientific inquiry must be kept separate from CLI. Moreover, their experiences also highlight the consideration that Asian and Asian American academics have with being seen as a representative of their race, hinting at the influence of the model minority discourse on their academic experiences. This contradiction between cultural connections spurring research questions but being siloed within the rest of the research process reveals how students develop sophisticated strategies for navigating institutional expectations.

### Navigational Capital: Strategic Compartmentalization

Our findings suggest that navigational capital—the skills to maneuver through institutions not created with communities of color in mind—underlies the “siloing” approach adopted by many participants. This strategic compartmentalization represents more than simple assimilation; it demonstrates how Asian science students have developed nuanced understandings of when and how CLI can be acknowledged within scientific institutions.

This navigation manifested most clearly in students’ ability to distinguish between different disciplinary and research contexts. When Nathaniel, an 18-year-old East Asian explained his approach to identity in research, he demonstrated sophisticated institutional navigation:

> I do data analysis, so I don’t really think it’s applicable…That would go into one of the bias categories for writing. I think if you’re doing a meta-analysis on your own culture or something like that, or your doctorate is something culture related, I think it’s fine to know your biases and stuff.

For Nathaniel, ethnic identities bias the scientific process and thus should be relegated to other disciplines. His distinction between contexts where acknowledging cultural background is “fine” versus where it represents “bias categories for writing” reveals a nuanced ability to read and respond to different perceived institutional contexts. In other words, he believed that calling out one’s CLI is irrelevant in scientific studies that are not focused on ethnic, racial, or linguistic identities.

Similarly, Evelyn, an 18-year-old East Asian woman, also demonstrated strategic compartmentalization when she siloed ethnic identity from her type of “basic science” research, stating:

> In my opinion it wasn’t necessary, at our point of study to kind of bring in more cultural or racial viewpoints. But I do definitely think, especially with, like, clinical studies or, you know, when you actually talk about like, how is this treatment going to be used, that viewpoint should definitely be considered. I just think with, like, since you’re doing more like drug discovery, and just on the more like basic science aspect, I didn’t express any, and I didn’t think it was kind of necessarily appropriate to do so.

The pattern extended to other participants who had internalized disciplinary hierarchies about where identity belongs. Abigail, a 17-year-old East Asian student, shared, “I mean, I don’t see myself bringing in my racial or cultural points in my research because I’m more into like biological sciences.” And Tanusha, a 18-year-old woman who identified as Southeast Asian, explained, “I don’t see the necessity of bringing that aspect into my writing, because it is primarily scientific…But I don’t think there’s like a need for me to like insert my own identity into like my project, because it is solely focused on like very molecular biology.” For these Asian and Asian American science students, then, their experience researching and publishing in the biological sciences was rendered as incompatible as being shaped by their ethnic identities, even if that type of connection might be allowed in other disciplines.

Whereas some Asian and Asian American science students made room for the incorporation of ethnic identities within the scientific process within non-STEM disciplines, others (73.9%) also explained that they didn’t feel that ethnic identities should ever make their way into scientific writing. For example, Nhan, a 20-year-old Southeast Asian man, said

> So, I could see why people might want to write and say, when you’re writing, uh, uh, an acknowledgement section, you could talk about yourself a little bit, I guess. But I don’t think it’s designed necessarily to accommodate a lot of story sharing, right? If you want to know more about somebody, you can go and read a biography. But if you want to know what scientific contributions they’ve made, you go and read their papers, right? You don’t go to science paper expecting, you know, gossip or anything like that.

And Peihua, an 18-year-old who identified as Pacific Islander and East Asian, expanded on this “I don’t think that there’s ever really much of an opportunity in this type of scientific writing, like in papers, um, to talk about your own identities, because like, It seems like you’re not supposed to talk about it.” For these student academic writers, then, publishing a scientific paper necessitated a kind of bifurcation between who they were as an Asian researcher and the process of science. These narratives show the recognition of implicit cultural scripts in science and reveal strategic identity management on the part of our interviewees. Moreover, these narratives also demonstrate how Asian science students sometimes perceive having to navigate institutional power dynamics and ethnic/ racial stereotypes to be perceived as legitimate scientists.

## Discussion

Our analysis reveals complex dynamics in how Asian and Asian American science students navigate cultural identity in scientific writing. Two primary approaches emerged: *spurring*, wherein cultural backgrounds served as catalysts for scientific inquiry, and *siloing*, wherein cultural identity was deliberately separated from scientific work. These complementary strategies reflect different ways of deploying cultural capital and navigating institutional expectations within scientific contexts. Students who engaged in spurring and siloing drew on both familial and navigational capital to empower them to engage in scientific inquiry and communication in ways that made sense to them. These approaches directly address our research questions regarding how Asian and Asian American science students draw upon cultural capital, view scientific writing as a space for identity integration, and respond to dominant scientific norms. Indeed, as people identify, access, and apply their capital, they often feel more agentic (Author et al., 2024).

Our findings suggest that these students strategically manage their cultural identities, drawing on them for motivation and research direction while often compartmentalizing them during the formal writing process. This navigation reveals both the potential contributions of diverse cultural perspectives as capital, supporting the use of translanguaging strategies in nuanced ways (Licona & Kelly, 2020). While students may not have engaged in explicit translanguaging (integrating discourses from their home language with academic science), they drew on ethnically relevant knowledge to engage in scientific inquiry. Students who were inspired by their cultural knowledge recognized the expanded capital on which they could draw as they conducted and communicated science. Students who separated these may or may not have seen their cultural capital as an asset but chose to not reference it in their scientific inquiry and writing process, a demonstration of their own navigational agency (Song, 2024).

While our data provide evidence for familial and navigational capital, we found limited direct evidence for other forms of cultural capital described by Yosso (2005), such as aspirational, linguistic, and resistant capital. For example, aspirational capital—the ability to maintain hopes and dreams despite barriers—might be implied in Evans statement that his background increased his passion for his work, but our data do not explicitly capture participants overcoming significant obstacles through cultural resilience. Similarly, while many participants likely possessed linguistic capital as multilingual individuals, our interviews did not specifically address how this might enhance their scientific communication. However, we acknowledge that our interviews were conducted in English, decreasing the opportunities for participants to leverage linguistic capital and reinforcing linguistic imperialism (Morrison & Lui, 2000).

Resistant capital—knowledge and skills fostered through challenging inequality—was not prominently featured in our data, possibly reflecting the early career stage of our participants or the constraints of scientific contexts that may discourage explicit resistance to dominant paradigms (Revelo & Barber, 2018). However, resistance may be demonstrated through actions and not through explicit acknowledgement of it during interviews. By positioning themselves as science writers and not Asian or Asian American science writers, the participants likely drew on cultural capital – the knowledge of how to reposition themselves within different communities of practice (Erol, 2010; Song, 2024). In addition, the students in this study were participating in research and writing activities that were often not expected in their academic classrooms; they were choosing to do more. Although, the act of being engaged in co-curricular science inquiry and writing may have been encouraged by their familial, academic, or cultural communities, we do not have explicit data indicating this. Therefore, we assume that the science students were agentic, regardless of whether they spurred or siloed their CLI from their academic science writing.

### Scientific Writing as a Site of Identity Navigation

Our findings reveal scientific writing as a complex site where Asian and Asian American science students strategically manage their identities in response to perceived norms and expectations. The “siloing” approach demonstrates sophisticated identity management, where participants consciously separate their cultural identities from their scientific work. The strategic compartmentalization appears particularly pronounced in certain disciplines. As Tanusha explained, science and CLI can be separate. Her perceptions reflect both an understanding of disciplinary norms and a strategic decision to conform to them. Similarly, Evelyn’s distinction between “basic science” research, where cultural viewpoints seemed inappropriate, and “clinical studies,” where they might be more relevant, shows nuanced navigation of disciplinary boundaries.

Most participants expressed that ethnic identities should not enter scientific writing, reflecting a deeply internalized norm of objectivity in science (Medin & Bang, 2014). Nhans statement that scientific papers are not intended to be stories when he contrasted biographies and scientific papers reflects this sharp division between CLI and academic/professional identity. Furthermore, our participants held strong beliefs in scientific objectivity and the corresponding view that incorporating cultural perspectives constitutes “bias.” This disciplinary variation aligns with Savolainen et al.’s (2023) observation that some fields view acknowledging positionality as violating scientific principles of universalism. Indeed, border crossing from academic to familial spaces, or translanguaging, involves knowing what, how, and when to communicate in certain ways to complete tasks successfully (Pun & Tai, 2021). The fact that several of the participants were aware of this strategy demonstrates their own agency in navigating diverse discourse spaces (Song, 2024).

The characterization of cultural perspectives as potential sources of “bias” rather than as valuable epistemological resources reveals a limited understanding of how all knowledge is situated. As feminist philosophers of science like Harding (1991) have argued, pretensions to pure objectivity often mask the ways that dominant perspectives shape scientific inquiry. Our participants’ concern with avoiding “bias” reflects what Medin and Bang (2014) describe as the myth of cultural neutrality in science—the mistaken belief that scientific knowledge exists independent of cultural frameworks. Some of our findings align with Chinn’s (2002) work on Asian and Pacific Islander women scientists, which revealed similar tensions in identity navigation. However, while Chinn’s participants often wished they “were males” to better fit scientific contexts, our participants instead chose to compartmentalize their cultural identities while maintaining them outside scientific contexts. This difference may reflect generational changes in how minoritized scientists manage identity tensions. The model minority stereotype likely influences these identity navigation strategies for our participants. As Lee (1996) documented, this stereotype creates expectations that Asian Americans will succeed academically while remaining culturally “other.” Our participants’ careful management (whether consciously or subconsciously) of cultural identity may represent a response to these contradictory expectations—demonstrating scientific competence while selectively controlling when and how cultural background becomes visible (Erel, 2010). Students may have been challenging and resisting cultural essentialism, for example (Song, 2024).

The commitment to scientific objectivity creates tension for students who draw upon cultural knowledge to identify research questions (our “spurring” findings) but feel compelled to remove traces of this influence from formal scientific writing. This contradiction allows Asian and Asian American students to reach scientific audiences with culturally motivated questions while obscuring the epistemological contributions their perspectives might offer. Our findings align with frameworks describing meaning-making across cultural boundaries (Aikenhead & Jegede, 1999; Moje at al, 2004), particularly Aikenhead’s concept of border crossing as students navigate between their home culture and science culture. Costa (1995) describes four different border transitions, ranging from “smooth” to “impossible”. In smooth transitions, students may easily cross between cultural and academic spaces with ease because there is such congruence between the spaces, this is compared to “hazardous” crossings, where stark cultural differences require more deliberate negotiation (Costa, 1995). For our predominantly U.S.-based participants (87%) the border crossing experience appears complex, but what Costa (1995) would refer to as “manageable.”

The evidence in our study suggests that border crossing is not simply a passive experience but an active, strategic process of cultural negotiation. Students engaging in in “spurring” demonstrated by what Moje and colleagues (2004) term “linking”—finding commonalities between cultural knowledge and academic discourse (such as Chloe’s familiarity with traditional medicine or Evan’s environmental concerns in Chinese communities) and academic discourse. This represents successful border crossing where cultural identity empowered scientific inquiry. However, the predominant pattern reflected systematic border crossing—keeping different knowledge systems separate—without integration (Aikenhead & Jegede, 1999). Students like Nathaniel and Evelyn carefully negotated which aspects of their cultural identity could cross into scientific spaces and which must remain on the “home” side of the border. Their strategic compartmentalization—determining where cultural identity seems “appropriate” versus where it might constitute “bias”--reveals sophisticated understanding of institutional boundaries.

Importantly, Aikenhead’s framework helps us understand “siloing” not merely as compartmentalization, but as a conscious secondary strategy when model minority narratives fail to advantage the construction of scientific identities. When students recognized that their Asian identity might mark them as biased, rather than bringing valuable perspective, they strategically chose to keep cultural and scientific knowledge separate. This suggests that the choice between spurring and siloing reflects ongoing cultural negotiation based on students’ assessment of how their identity will be perceived in specific contexts.

The post-immigration context of our participants adds another layer to border crossing dynamics. Most participants identified as from the United States, meaning their “home culture” is itself likely a hybrid of Asian heritage and American socialization. This may explain why some students initially reported not thinking about their culture in their work. Their everyday border crossings between home and school may have been relatively smooth until they encountered the specific culture of scientific writing, with its emphasis on objectivity. The model minority stereotype may have initially facilitated smooth crossings by positioning them as naturally suited for science, only to create hazardous borders later when they attempted to explicitly consider cultural perspectives. As the majority of participants indicated, they learned that ethnic identities should never enter scientific writing, a clear message about which borders cannot be crossed and what must remain siloed.

Notably absent from our findings was evidence of blending - the integration of different knowledge systems that has been described as the most sophisticated form of meaning-making (Moje et al., 2004). Blending as a form of translanguaging can allow students to feel motivated as they acquire new knowledge while referencing what is familiar (Rafi & Morgan, 2023).The institutional pressures our participants described suggest that scientific writing contexts may actively discourage blending, pushing students toward border crossing as the safest navigational strategy. As Sanchez Tapia (2020) noted, not all students may use different capital during academic activities if they have learned that cultural knowledge should be maintained separately from academic contexts. Our participants’ strategic compartmentalization reflects precisely this learned separation, suggesting that institutional norms may systematically prevent the blending that could enrich scientific inquiry. Song (2024) found that Chinese American students challenged the expectation of using language to satisfy economic interests and maintain cultural cohesion.

### Theoretical Implications

Our findings extend CCW theory by demonstrating how cultural capital operates in scientific contexts that may not readily acknowledge its value. While Yosso (2005) developed this framework primarily for educational settings, our research shows how it applies to professional scientific practice, particularly scientific writing and publication. Importantly, our findings reveal that cultural capital can operate even when not explicitly acknowledged—driving research questions and motivation even when compartmentalized in formal scientific writing. Similar discussions of the strategic management of social identities have documented the ways in which people from marginalized groups may deploy their identities in professional contexts (Erskine, Bonner, & Rabelo, 2025; Roberts, Cha, & Kim 2014; Roberts, Settles, & Jellison, 2008). The complementary “spurring” and “siloing” strategies we identified contribute to understanding strategic identity management in scientific writing contexts Unlike binary models that position scientists as either assimilating to or resisting dominant norms, our findings suggest a more nuanced process where CLI is selectively deployed at different stages of the inquiry process (Pun & Tai, 2021). This selective deployment allows science students to both benefit from cultural resources and conform to institutional expectations. Hence, the participants in our study found how to successfully engage in border crossing.

Taking a critical approach to Yosso’s framework based on the recently published work of Song (2024), we may view “spurring” in a different light. While spurring may demonstrate an activation of cultural capital, using a framework from Song we could also interpret this as “bargaining” - attempting to make their cultural resources legible and valuable within scientific institutions. Similarly, in the same lens, we may view “siloing” as students’ recognition that their cultural resources lack institutional recognition within scientific contexts, leading them to compartmentalize rather than risk devaluation. This inherently calls to question who gives capital to culture? In this instance of academic writing, the publishers, editors, and reviewers deem what is acceptable and therefore what culture has capital. Our work highlights the work of others (Ramírez-Castañeda, 2020) advocating for more inclusive publishing practices that explicitly welcome the multiple ways academic research may be expressed.

Our research also contributes to literature that challenges the model minority discourse by showing that science students did not conflate their scientific achievement with their ethnic/cultural identities (Poon et al., 2016). The perception that acknowledging cultural influence constitutes “bias” reflects the model minority expectation that Asian Americans will succeed by adopting dominant norms rather than challenging them. However, our participants’ strategic navigation of when and how to incorporate cultural perspectives demonstrates agency within these constraints. Our findings allow us to extend Erel’s (2010) findings that as migrants move across cultural spaces, they must contend with intra-cultural issues of validation, such as knowing which languages afford power in certain environments. Likewise, as Asian and Asian American science students move back and forth from academic to familial spaces (i.e., engage in border crossing), they likely learn how to identify and leverage relevant capitals.

### Implications for science education

Our findings suggest important implications for science education. First, educators should help students recognize how cultural backgrounds can serve as valuable resources rather than biasing factors in scientific inquiry. By explicitly acknowledging the role of cultural perspectives in shaping research questions and approaches, educators can help students leverage their cultural capital more effectively. Second, science education should address the myth of pure objectivity, helping students understand that all knowledge is situated in cultural contexts. Teaching about the history and philosophy of science could help students develop more nuanced understandings of objectivity that recognize the value of diverse perspectives rather than attempting to erase them (Author et al., 2024). Finally, science educators should provide models of successful identity integration in scientific work. Exposing students to scientists who effectively bridge cultural knowledge with scientific practice could help them envision possibilities beyond the binary choice between cultural authenticity and scientific legitimacy.

Scientific publication venues might consider how their formats and expectations could evolve to better accommodate diverse backgrounds. While maintaining rigor, journals could explore alternative formats that allow for more explicit acknowledgment of researchers’ positionality and the cultural contexts that inform their work. Additionally, peer review processes could be developed to recognize the value of diverse cultural perspectives rather than treating them as potential sources of bias. Reviewers might be trained to distinguish between methodological rigor and cultural neutrality, acknowledging that good science can emerge from culturally situated perspectives.

### Limitations and Future Research

Our study has limitations that suggest directions for future research. Our sample focused on Asian and Asian American students, limiting generalizability to other ethnic/ethnic groups and career stages. Future studies could explore how scientists from other minoritized groups navigate identity in scientific writing, as well as how these strategies evolve throughout scientific careers. While our interview protocol was designed to specifically target certain forms of cultural capital such as resistant and linguistic capital, participants seemed reluctant to share or did not see these questions as relevant to their science research and writing, despite both interviewers being near peers of Asian descent. We also did not collect and analyze data regarding the participants’ science mentors. Future studies could more deliberately explore how science students draw upon aspirational capital to persist through challenges, leverage multilingual abilities in scientific communication, or develop ways to respectfully challenge dominant paradigms. Additionally, educational interventions could help students recognize these forms of capital as potential resources rather than viewing cultural backgrounds as irrelevant or potentially biasing in scientific contexts. Future research could more deliberately explore these aspects through targeted questions about institutional navigation and multilingual communication strategies. Future work can also explore whether such navigational capital, as we observed as outsiders, is intentional or perceived as necessary on the part of the science students themselves. Longitudinal studies would be particularly valuable for understanding how identity navigation strategies develop over time. Following scientists from training through career advancement could reveal how institutional contexts and career pressures shape identity expression in scientific writing over time and professional experience.

## Conclusion

This study reveals both the potential contributions of diverse cultural perspectives to scientific knowledge and the persistent constraints that limit their full expression. By documenting how Asian science students strategically navigate between cultural identities and scientific norms, we contribute to understanding both individual agency and institutional constraints in scientific practice. The complementary strategies of “spurring” and “siloing” demonstrate sophisticated identity management that allows scientists to draw upon cultural resources while meeting institutional expectations. These strategies represent not just personal choices but responses to broader scientific norms and ethnic dynamics.

## Ethics Statement

The study was approved by [x] University Institutional Review Board ([study number]. All participants provided their informed consent or assent prior to completing the interview.

## Statements and Declarations

## Ethics and Consent to Participate

The study was approved by the Emory University Institutional Review Board (STUDY00000797). All participants provided their informed consent prior to completing the interview.

## Data availability

Anonymized data available upon reasonable request

## Authors Contributions

DG: Project ideation, data collection, data analysis, manuscript writing and editing

SS: data collection, analysis, manuscript writing and editing

GK: data collection, analysis, manuscript writing and editing

LTS: data collection, manuscript editing

MB: manuscript writing and editing

SCF: Project ideation, data collection, data analysis, manuscript writing and editing

## Biographical Note

Devon Goss is an Associate Professor of Sociology at Oxford College of Emory University sociologist specializing in the areas of race and ethnicity, the family, education, and sports. She completed her PhD in 2018 at the University of Connecticut. Much of her research centers on the experiences of those who cross racial boundaries. Her co-authored book with SunAh Laybourn, *Diversity in Black Greek-Letter Organizations: Breaking the Line* (2018), explored non-Black members of African American sororities and fraternities. Previously, she documented the impact of racial ideology on members of transracial families, the experiences of white students who attend Historically Black Colleges and Universities, the racial motivations of white expats in Mexico, and the racialization of gymnastics sports commentary. Her current research project explores the ways that colleges and universities narrativize their histories with enslavement.

Shaila Sachdev is an undergraduate student at Emory University, majoring in Biology and Predictive Health. She is passionate about promoting science literacy and making scientific research more accessible to broader audiences through involvement in student-led research publications and courses focused on the evolution of the scientific research article. She is also deeply committed to health education and currently volunteers with an organization that mentors high school students aspiring to enter healthcare and science fields.

Grace Kim spent her first two years of undergraduate studies at Oxford College of Emory University where she took classes in Sociology of Education and tutored students in mathematics. She is interested in education equity and is currently volunteering at a nonprofit organization to help low income, high achieving high school students navigate the college application process. She is now at Dartmouth College studying mathematics.

LaTonia Taliaferro-Smith is an Associate Teaching Professor of Biology at Oxford College of Emory University. She actively engages undergraduate students in her research investigating triple negative breast cancer. She is also interested in the intersection of race, ethnicity, and cancer biology.

Meena Balgopal is a Professor in the Department of Biology and a University Distinguished Teaching Scholar at Colorado State University. Her research group explores how people make meaning of natural science concepts with a focus on writing, reading, quantitative reasoning, and place-based education. She is also examines how disciplinary literacies shape students’ academic identities and motivations to remain in the sciences.

Sarah C. Fankhauser is an Associate Professor of Biology at Oxford College of Emory University. While her academic training is in microbiology, she transitioned her research focus many years ago to examine how authentic disciplinary literacy practices impact students’ identities, motivations, and critical scientific literacy.

